# Jointly Modeling Species Niche and Phylogenetic Model in a Bayesian Hierarchical Framework

**DOI:** 10.1101/2022.07.06.499056

**Authors:** Sean W McHugh, Anahí Espíndola, Emma White, Josef Uyeda

**Affiliations:** Biological Sciences, Virginia Tech; Department of Entomology, Plant Sciences Building 3138, 4291 Fieldhouse Dr., University of Maryland, College Park, MD 20742-4454, USA

## Abstract

When studying how species will respond to climatic change, a common goal is to predict how species distributions change through time. Environmental niche models (ENMs) are commonly used to estimate a species’ environmental niche from observed patterns of occurrence and environmental predictors. However, species distributions are often shaped by non-environmental factors–including biotic interactions and dispersal barriers—truncating niche estimates. Though a truncated niche estimate may accurately predict present-day species distribution within the sampled area, this accuracy decreases when predicting occurrence at different places and under different environmental conditions. Modeling niche in a phylogenetic framework leverages a clade’s shared evolutionary history to pull species estimates closer towards phylogenetic conserved values and farther away from species specific biases. We propose a new Bayesian model of phylogenetic niche estimation implemented in R called *BePhyNE* (Bayesian environmental Phylogenetic Niche Estimation). Under our model, species ENM parameters are transformed into biologically interpretable continuous parameters of environmental niche optimum, breadth, and tolerance evolving as a multivariate Brownian motion. Through simulation analyses, we demonstrate model accuracy and precision that improve as phylogeny size increases. We also demonstrate our model on eastern United States Plethodontid salamanders and recover accurate estimates of species niche, even when species occurrence data is lacking and entirely informed by the evolutionary model. Our model demonstrates a novel framework where niche changes can be studied forwards and backwards through time to understand ancestral ranges, patterns of environmental specialization, and estimate niches of data-deficient species.

## I. INTRODUCTION

### 1. Summary of Current Methods to Model Niche Evolution

The biotic and abiotic environment shapes species’ ecological niches, often determining a species’ past and future chances of survival (Losos 2008; Evans et al. 2009; Ogburn and Edwards 2015; Alves et al. 2017; Farallo et al. 2020). Understanding the macroevolutionary history of niche evolution is therefore critical to understanding past, current, and future biodiversity patterns. In this context, our ability to model the evolution of ecological niches can have conceptual but also practical consequences, since it can allow us to understand macroevolutionary constraints and patterns in the evolution of these important traits, as well as predict likely biodiversity changes under future environmental conditions or improve niche estimates for species for which few observations are available (e.g., rare species). While several recently-developed methods have tackled this challenge, researchers generally use two-stage analyses, in which species’ niches are first estimated and then analyzed in a comparative framework (Graham et al. 2004; Knouft et al. 2006; Warren et al. 2008; Evans et al. 2009; Heibl et al. 2018; Guillory and Brown 2021; Quintero et al. 2022). While such approaches can help uncover evolutionary patterns of species niche (Kozak and Wiens 2016; Kolanowska et al. 2017; Farallo et al. 2020; Gaynor et al. 2020), they result in models where present-day niche estimates inform evolutionary processes, but not the reverse. Furthermore, this approach can exacerbate biases, as closely-related species with competitive exclusion but apparently extensive access to suitable habitat may appear more divergent than they actually are (Owens et al. 2013; Saupe et al. 2018).

A common approach used to estimate and delimit species’ ecological niches is offered by Ecological Niche Models (ENMs), which most of the time estimate niches from current occurrence data and environmental (often climatic) predictors (Warren et al. 2008, 2010; Cooper et al. 2010; Smith et al. 2019). This method allows understanding how a species abiotic (i.e., fundamental, *sensu* Hutchinson, 1957) niche and eventually species ranges may change in response to environmental change (Quintero and Wiens 2013; Saupe et al. 2014; Ogburn and Edwards 2015; Farallo et al. 2020). However, such abiotic niches -and their spatial projections-exclude by definition the effect of biotic factors and dispersal barriers that can reduce ranges to a subsection of the actual extent of the abiotic niche (Soberón 2007; Boulangeat et al. 2012). In such situations, these models can result in biologically-unrealistic species ranges, especially under projected climate scenarios (Elith and Leathwick 2009; Peterson and Soberón 2012; Huang and Frimpong 2015). Further, ENMs generally ignore evolutionary processes and information on the niches of related species (Franklin and Miller 2009; Blank and Blaustein 2012; Oke et al. 2014), which can also affect the quality of the estimates and their reflection of the natural and evolutionary history of the species or clade in question. Finally, restricting niche estimates to extant occurrence data can give a skewed view of species niches and their evolutionary dynamics, especially given recent climatic and anthropogenic change.

Phylogenetic models of niche evolution hold potential to improve both species niche estimates and our understanding of their evolution. Such methods would ideally be tied to biologically-meaningful and interpretable parameters that could be connected to, for example, physiological tolerances of organisms. Today, a handful of approaches allow the inclusion of phylogenetic structure in niche estimation using linear mixed modeling (Hadfield and Nakagawa 2010; Ives and Helmus 2011; Morales-Castilla et al. 2017; Li et al. 2020; but see Quintero et al. 2022). For example, phylogenetic information can be integrated through the use of phylogenetic predictors (Morales-Castilla et al. 2017) or phylogenetically-covarying random effects (Ives and Helmus 2011). However, in those cases, the estimated parameters are not easily tied to meaningful and biologically-interpretable evolving traits (e.g., logistic regression coefficients cannot be interpreted in isolation, restricting their utility in directly modeling different aspects of niche under biologically-meaningful evolutionary processes). Other approaches, such as that of Hua et al. (2021), address this problem through a sophisticated event-based model that fits from root to tips of the phylogeny a history of adaptation, dispersal, and speciation that best explains the observed variation in niche predictors. While such reconstructions can model complex mechanistic niche history into a composite of repeated, predefined niche-shaping events, the resulting reconstructions are hard to generalize, and largely disconnected from the large family of continuous models of stochastic trait evolution that are commonly used in phylogenetic comparative methods (Felsenstein 1985; Blomberg et al. 2003; Butler and King 2004; Beaulieu et al. 2012; Landis et al. 2013; Drury et al. 2016). These models provide flexible hypothesis-testing tools to compare evolutionary scenarios (O’Meara et al. 2006; Revell and Harmon 2008; Clavel et al. 2015; Adams and Collyer 2018). Thus, although much progress has been made, linking macroevolutionary models with biologically-meaningful parameters (e.g., niche optimum, breadth, and tolerance) is still needed. Doing this would allow us to more explicitly model how different components of the niche respond to changing environments (Condamine et al. 2013; Quintero and Wiens 2013; Qiao et al. 2016).

To address this methodological gap, we propose an alternative modeling framework in which ENMs and the evolutionary process of biologically-realistic environmental niche traits are modeled jointly. Here, we jointly model (1) current environmental responses of taxa in a clade from discrete occurrence observations, and (2) the continuous evolutionary processes shaping these environmental responses under a multivariate Brownian Motion random walk. Although we implement a multivariate Brownian Motion model (Felsenstein 1985, Figure 1), our approach could be readily expanded to include other models commonly used in phylogenetic comparative methods (e.g. Hansen 1997; Blomberg et al. 2003; Butler and King 2004; Beaulieu et al. 2012; Landis et al. 2013; Drury et al. 2016). We implement our model in the R package *BePhyNE* (R Core Team 2021).

**Figure 1:**
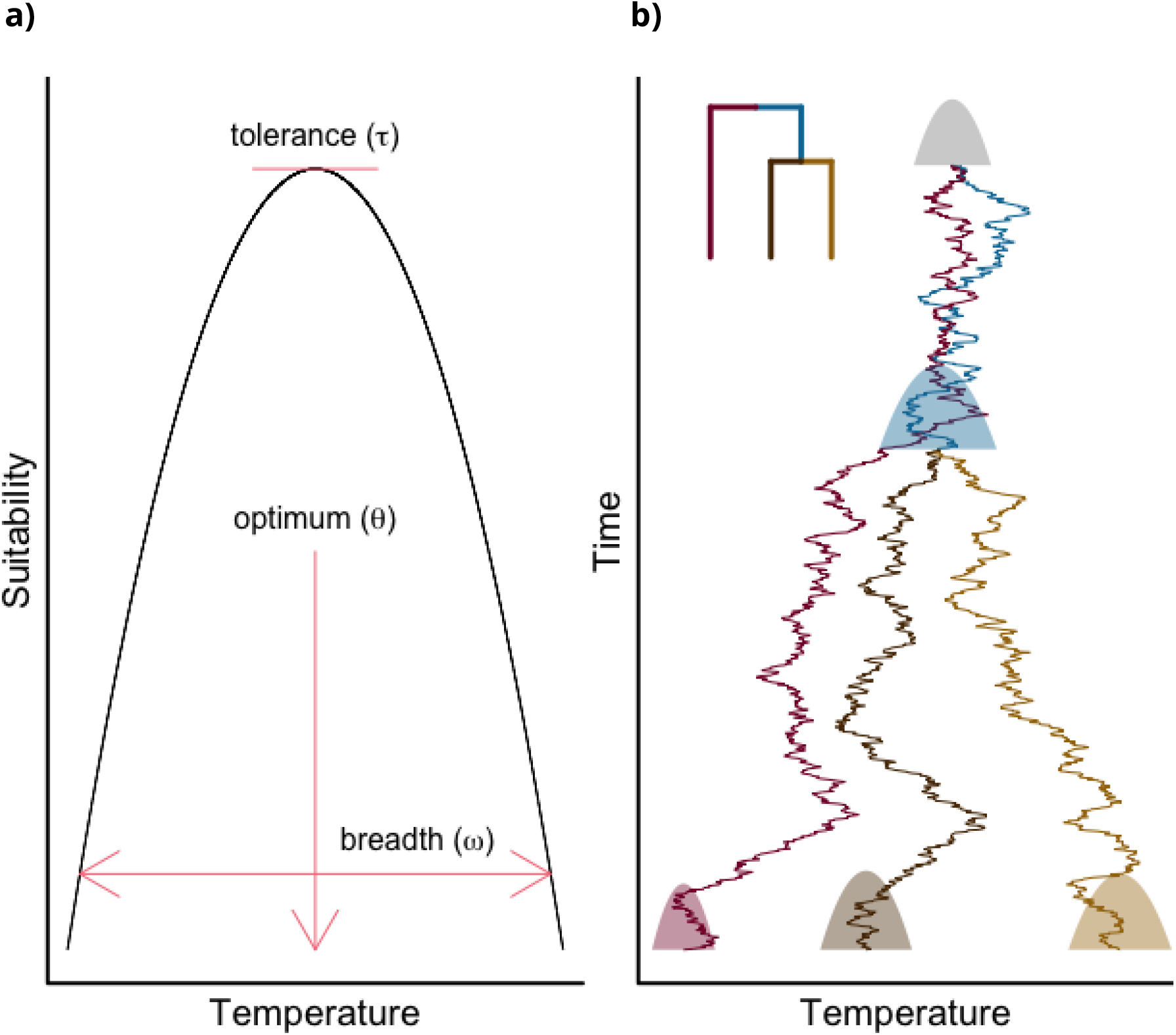
Environmental response curves at the species level and phylogenetic level are jointly modeled in *BePhyNE*. a) Species level response curves for continuous environmental predictors. Each species has three continuous characters describing its niche: an optimum (*θ*) where the species occurs at the highest probability, a breadth (ω) depicting the environmental range and limits for a species to occur at any probability above given threshold (.05), and a tolerance (*t*) indicating the highest probability the species has of occurring anywhere. b) At the phylogenetic level, macroevolutionary divergence in the environmental response parameters *θ*, ω, and *t* evolve under a model of stochastic multivariate continuous evolution as simulated over a three taxa tree (evolutionary history of *θ* is illustrated through time). Response curves are plotted at the root, nodes, and tips of the phylogeny.

## II: Model Description

### 1. Species-Level Niche

We developed a Bayesian framework for phylogenetic niche modeling in the statistical programming language R (R Core Team 2021) as an open-source package, *BePhyNE* (Bayesian environmental Phylogenetic Niche Estimation). We first consider the Grinnellian niche for a single measured continuous environmental predictor *x* (e.g., temperature, precipitation), where a species’ occurrence probability (*P_i_*, for species species *i*) is a function of the environmental predictor *x* (Fig. 2a) (Grinnell 1917; Soberón 2007). Specifically, a quadratic logit-linked generalized linear model (GLM) fits a curve (hereafter, “the response curve”) that relates the occurrence probability to the observed values of the predictor (Fig. 2b) (Braak and Looman 1986; Jamil and Braak 2013; Jamil et al. 2014). This GLM is then fit to binary presence/absence occurrence data for species *i, Y_i_*, such that:

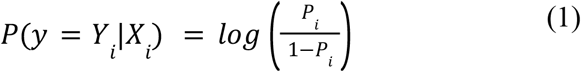

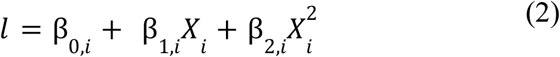

where *l* is the log-odds of observing species *i* given the measured environmental predictor data, *X_i_*, and the Bernoulli-distributed observations *Y_i_*. The vector of estimated regression coefficients, ***β***, describes the relationship between predictor values and the probability of occurrence. Unconstrained GLM’s fit to empirical data often estimate biologically-unrealistic response curves when occurrence data only cover a fraction of the abiotic niche space. For example, an estimate may not have an intermediate optimum for *x*, but rather an intermediate valley with occurrences having highest probability at the infinite extremes of the predictor. The interdependence of the ***β*** coefficients in equations (1) and (2) hinders modeling the evolution of ecological niche, as it requires a complex set of constraints to ensure the unimodality expected under most biologically-realistic scenarios. To remedy this, we reparametrize the model to constrain the response curve as a positive unimodal distribution. We define the unimodal niche response curve for predictor *j* in species *i* as a set of three parameters (Θ_ij_={*t_ij_, θ_ij_, ω_ij_*}). Each parameter has a direct biological interpretation, with values upon which we can set meaningful constraints: tolerance (a species highest probability of occurrence, *t*), optimum (predictor value at a species highest probability of occurrence, *θ*), and breadth (range of predictor values a species can occur at, *ω*) (Figs. 1a and 2c). These parameters are algebraically transformed to the GLM (***β***) coefficients, deterministically setting the GLM response curve (see Appendix I for the full derivation). The niche response curve parameters are linked to the underlying GLM regression coefficients (***β***) as:

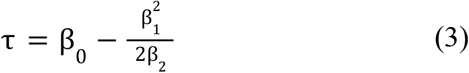

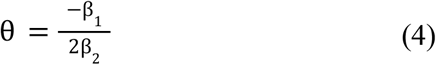

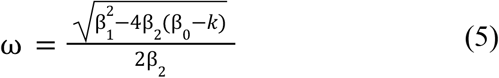

**Figure 2:**
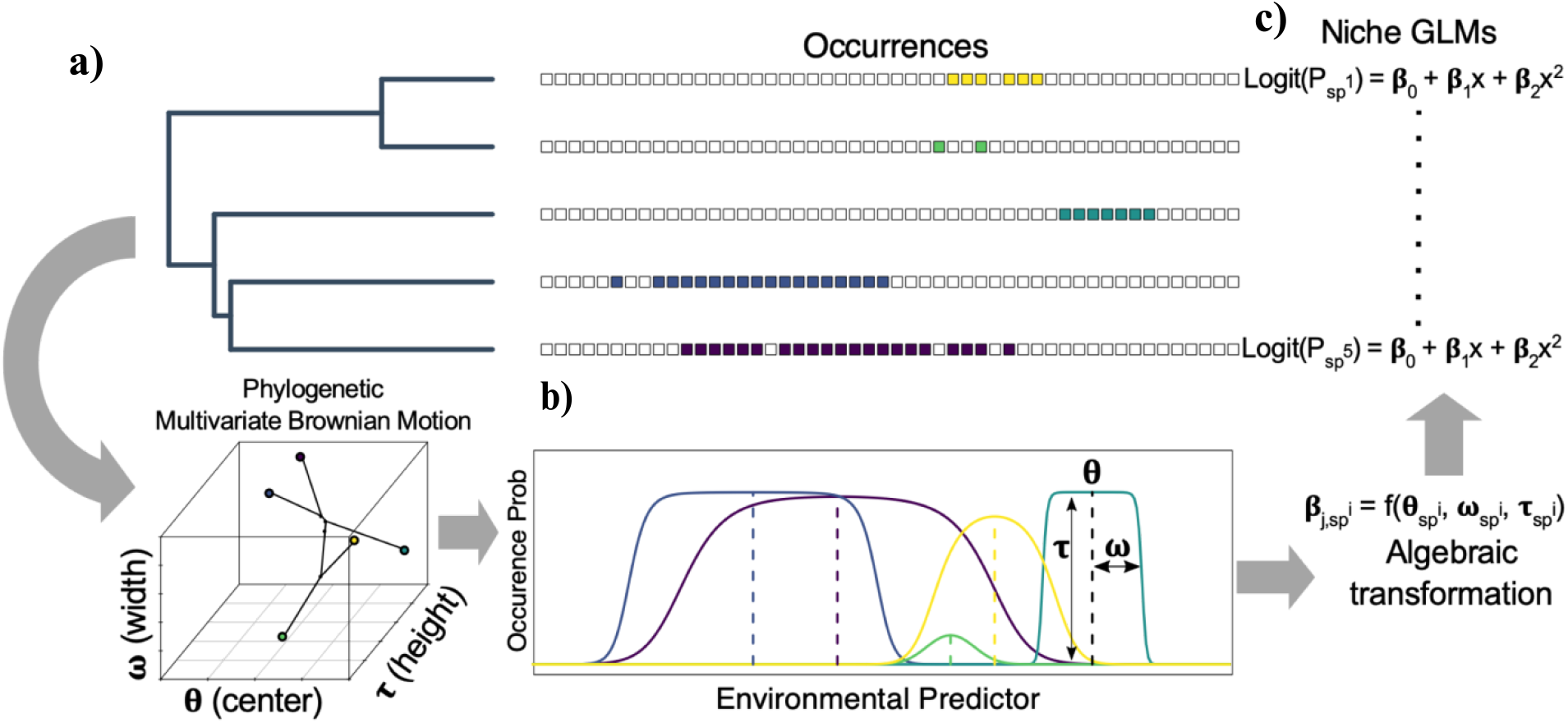
Bayesian estimation of the phylogenetic niche model used in this study, which allows for the joint estimation of species environmental response curves and their evolution using an algebraic link between species’ niche parameters and GLM coefficients. a) The phylogeny is used to model the joint evolution of the parameters describing the niche optimum (*θ*), breadth, (ω), and tolerance (*t*) for the b) unimodal response curves under a multivariate Brownian Motion model. c) For each species, we algebraically transform niche parameters into GLM coefficients (***β***), which are used to calculate the likelihood of the observed occurrence data. Throughout the Bayesian MCMC, we update species-specific niche parameters, as well as parameters governing the multivariate Brownian Motion model.

The parameter *k* represents a threshold cutoff probability across which the breadth is measured. The resulting set of niche parameters across all predictors and species in the phylogeny is defined as Θ_niche_.

### 2. Phylogenetic Niche Evolution

We assume that each of the three niche parameters in Θ_ij_ evolve over time on the phylogeny under a multivariate Gaussian process. Here, we implement a simple multivariate Brownian motion process (Felsenstein 1985, 2004; Revell et al. 2008) where parameters randomly diffuse through trait space along the phylogeny, determined only by a vector of phylogenetic root means (***A***) and the evolutionary rate matrix (***R***) describing the evolutionary rates and covariances of the three niche parameters (Figs. 1b and 2d). We decompose ***R*** into a vector of standard deviations (***σ***) representing the evolutionary rates, and a correlation matrix (*R_COR_*) that describes the evolutionary correlation between different parameters of a species’ response curve (Caetano and Harmon 2019). We denote the set of parameters defining the evolutionary model as *Θ_BM_* = {*A, R*}.

To simplify phylogenetic niche estimation over multiple environmental predictors, we assume the environmental predictors were evolutionarily uncorrelated. We assume each predictor *j* evolves under an independent mvBM process with an evolutionary rate matrix (*R_j_*) for *θ_j_* and ω_j_ of the response curve. While *θ* and ω vary independently across environmental predictors, the *t* parameter is necessarily linked to produce a single maximum probability of occurrence across environmental predictors. The biological interpretation of *t* is dependent on the occurrence data used. When using real presence-absence data, *t* is the maximum species occurrence probability–a biological trait that could reasonably be assumed to be phylogenetically heritable and potentially fit a Brownian Motion process. However, generating real absences data is often prohibitively labor-intensive and frequently substituted with randomly sampled background points or pseudo-generated absences. With pseudo-absence data, species response curves model relative suitability across environmental predictor values, changing the interpretation of *t* to one removed from the biologically-meaningful measures of occupancy or occurrence probability (Elith et al. 2006; Li et al. 2011; Peterson and Soberón 2012). To avoid biasing the more biologically-realistic *θ* and ω when using generated absences, we explored modeling *t* at the species level with 1) species-specific priors or 2) *t* fixed across all species. For simplicity, in our model performance simulations, we fixed *t* to a single value for all species.

Our hierarchical Bayesian model estimates species niche and niche evolution jointly over the array of environmental predictors X. The joint probability of the model is as follows:

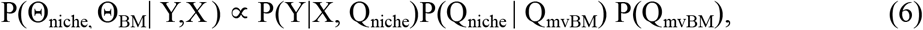

where P(Y|X, Q_niche_) is the probability of occurrence for each species given the predictor X and response curve parameters under the reparameterized GLM, P(Q_niche_ | Q_mvBM_) is the probability of the response curve parameters given the mvBM evolutionary process model, and P(Q_mvBM_) is the prior on the parameters of the evolutionary process.

### 3. mvBM Priors

Prior knowledge on species niche evolution is likely to vary greatly depending on the study system. Therefore, we implemented both informative and uninformative prior distributions on model parameters. For our informative priors, we implemented log-normal distributions for continuous parameters bounded by zero, including mvBM parameters: *A*_ω_, and ***σ***. Because *A_θ_* is continuous and unbounded, we implemented a normal distribution prior. For the *R_COR_* sub matrix, we implement an inverse-Wishart prior centered on a diagonal correlation matrix, where one can specify how strongly the prior is centered on a specified matrix by setting the degrees of freedom. For the inverse-Wishart distribution, an increase in the degrees of freedom will correspondingly result in a narrower distribution around the centered matrix. We specified weakly-informative diagonal matrices that assume no prior information on the direction of covariation, for example, between *θ* and ω.

### 4. MCMC Proposal Distributions

We estimated parameters of the model in the R statistical environment using Markov-Chain Monte-Carlo (MCMC) with a Metropolis-Hastings algorithm that stochastically explores parameter space (Chib and Greenberg 1995). We used efficient proposal and likelihood functions for the mvBM model of niche evolution from the package *ratematrix* (Caetano and Harmon 2017). *ratematrix* implements a separation strategy when proposing on *R*, where the matrix is decomposed into ***σ***, and *R_COR_*, with separate proposal and prior distributions for each (Barnard et al. 2000; Zhang et al. 2006; Liu et al. 2016). Species niche parameters and ***σ*** parameters are updated using both multiplier and sliding-window proposal distributions, while *A* are updated only using a sliding-window proposal. To limit proposals to biologically-realistic response curves, breadth is log-transformed to be constrained to positive trait space, while *t* is transformed to be on a 0.05-1.00 scale through a logit transformation to reflect the range of possible maximum occurrence probabilities for a species. The acceptance ratio for parameter proposals is:

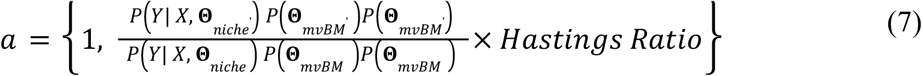

Terms in the numerator denote the probability under the proposal while terms in the denominator denote the probability under the current MCMC parameters. 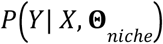 is the probability of occurrence given environmental predictor X and niche response curve parameters, 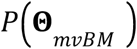 is the probability of the niche response curve across all species given the evolutionary process. 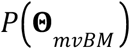 is the joint prior on the mvBM process parameters. Proposals are balanced by the Hastings ratios.

## III. Materials and Methods

### 1. Simulation Testing

We tested the accuracy, precision, and consistency of *BePhyNE* by simulating 500 datasets under the model and recovering the data-generating parameters (see Supplementary Materials 2 for description of additional test from Cook (2006)). We simulated and estimated parameters over 500 datasets sampled from the prior using 1 million generation MCMC at 10% burnin. We plotted the “true” parameters used to generate each simulated dataset against the posterior medians of the MCMC estimating over the same dataset. First, we sampled mvBM parameters from the prior distributions, to simulate response curve parameters (*θ* and *ω*) over a phylogeny simulated from a pure birth process using the R package *phytools* (Revell 2012). Niche tolerance *t* was fixed to 0.87 for all species for all predictors. Next, we transformed the response curve parameters for each tip to GLM coefficients, ***β***, which were applied across a simulated grid of virtual environmental predictor values to generate the occurrence probability (using Eq. (1)) at each grid cell. Given the cell’s occurrence probability, we sampled from a Bernoulli distribution to categorize the cell as a presence or absence. Finally, we fit our model to the simulated data to recover the “true” data generating parameters. We compared the simulated parameter values to MCMC posterior medians. We expect there to be a 1:1 linear relationship between the “true” and posterior median values. We assessed model consistency by testing if the accuracy and precision of model parameter estimates improved over datasets simulated from phylogenies with more tips. To do so, we repeated our larger simulation analyses over three treatments of 50 datasets simulated from prior distributions with 20, 50 and 100 phylogenetic tips, and compared accuracy and precision in parameter estimation across treatments.

### 2. Empirical Application: Eastern North American Plethodontidae

We fit *BePhyNE* to empirical occurrence data available for 82 species from the family Plethodontidae used in previous studies (Kozak and Wiens 2010b; Fisher-Reid et al. 2012). We matched the data to a larger 516-species time-calibrated phylogeny estimated by Bonett and Blair (2017) and pruned out data for missing species. We collected Plethodontidae occurrence data from online databases using a pipeline script coded in R. We used the R package *rgbif* (Chamberlain et al. 2014) to download occurrence data from the Global Biodiversity Information Facility (GBIF, (April 2021; ww.gbif.org)) into the R environment. We filtered out occurrences with missing or inaccurate spatial information using the package *CoordinateCleaner* (Zizka et al. 2019). Using the R package *dismo* (Hijmans et al. 2020), we generated target group pseudo-absences for each species from the presence data of other species. We downsampled occurrences so that presence/absence prevalences within a species expert range (as identified by IUCN expert range maps) were 50% (Mateo et al. 2010; Barbet-Massin et al. 2012). We added additional target absences equal to the number of absences within the expert range outside of it to include sampling across a wider abiotic range. For all occurrence and background points, we collected climatic data from the Worldclim v2.0 database on mean annual precipitation (mm) and mean annual temperature (°C) (Fick and Hijmans 2017). We divided our occurrence dataset in half to create a training set to estimate parameters on, and a validation set to predict species ranges from our parameter estimates.We chose empirical priors for ***A*** and ***σ*** in scaled parameter space based on the ranges of values of predictors from occurrence data and exploratory performance across a range of simulation analyses (Table 1). For example, the prior for *A_θ_* was set to a normal distribution with mean and standard deviation equal to the environmental averages and variation over the entire sampled area (including presences and absences). Evolutionary rate priors generate a range of response curves ranging from nearly complete stasis to complete saturation of the scaled space. For evolutionary correlations, we used an inverse-Wishart prior on *R_COR_* centered on a diagonal (uncorrelated) matrix with degrees of freedom equal to the diagonal matrix row length plus one—3.0.

**Table 1.** *BePhyNE* mvBM parameter priors and estimates for Plethodontid salamanders. *A:* parameters for the root means; *σ*: evolutionary rate parameters for each continuous trait indicated by the subscript (either niche optimum *θ* for or niche breadth *ω*); *R_COR_* : correlations between *θ* and *ω* for each environmental predictor.

| Parameter | Prior Distribution | Prior Distribution Parameters unscaled | Prior Distribution Parameters scaled | Posterior Median | 95% Highest Posterior Density |
| --- | --- | --- | --- | --- | --- |
| mean annual precipitation<br>$A_\theta$ (mm) | normal | mean=126.6<br>sd=13.1 | mean=0<br>sd=0.5 | 137.24 | 120.16–154.85 |
| mean annual precipitation<br>$A_\omega$ (mm) | log-normal | log-mean=7.9<br>sd=5.2 | log-mean=0.3<br>sd=0.2 | 9.94 | 6.82–13.81 |
| mean annual precipitation<br>$\sigma_\theta$ ($\frac{mm}{million\ years^{\frac{1}{2}}}$) | log-normal | log-mean=5.3<br>sd=29.1 | log-mean=0.2<br>sd=0.46 | 29.47 | 26.82–32.20 |
| mean annual precipitation<br>$\sigma_\omega$ ($\frac{mm}{million\ years^{\frac{1}{2}}}$) | log-normal | log-mean=12.0<br>sd=29.1 | log-mean=0.2<br>sd=1 | 17.94 | 15.33–20.67 |
| mean annual precipitation<br>$R_{COR}$ | Inverse-Wishart | df=3<br>centered<br>matrix=diagonal matrix | — | 0.10 | -0.01–0.21 |
| mean annual temperature<br>$A_\theta(^{\circ}\text{C})$ | normal | mean=11.7<br>sd=1.79 | mean=0<br>sd=0.5 | 12.2 | 9.63–14.7 |
| mean annual temperature<br>$A_\omega(^{\circ}\text{C})$ | log-normal | log-mean=1.1<br>sd=0.7 | log-mean=0.3<br>sd=0.2 | 1.70 | 1.17–2.35 |
| mean annual temperature<br>$\frac{\sigma_\theta(^{\circ}\text{C})}{\text{million years}^{\frac{1}{2}}}$ | log-normal | log-mean=0.7<br>sd=9.0 | log-mean=0.2<br>sd=0.46 | 3.47 | 3.17–3.78 |
| mean annual temperature<br>$\frac{\sigma_\omega(^{\circ}\text{C})}{\text{million years}^{\frac{1}{2}}}$ | log-normal | log-mean=1.6<br>sd=4.0 | log-mean=0.2<br>sd=1 | 2.94 | 2.59–3.33 |
| mean annual temperature<br>$R_{COR}$ | Inverse-Wishart | df=3<br>centered<br>matrix=diagonal matrix | — | -0.15 | -0.25 – -0.05 |

We ran 10 chains for 5 million generations with 20% burnin to assess convergence. We adjusted tuning parameters to approximate parameter acceptance ratios of 20%-40%. Using the validation dataset, we assessed the predictive performance of our species response curve estimates. We predicted the probability of occurrence for each species at each site in the validation dataset. We compared predicted occurrence probabilities to the real observed state of the site using a receiving operator characteristic (ROC) curve, a performance measure for binary classifiers. The ROC curve is a plot of true positive rates over false positive rates in predicted occurrence probabilities at probability thresholds ranging from 0 to 1 (Jiménez-Valverde 2012). Accurate response curve estimates display high true and low false positivity rates ratios, with the highest ratio approaching a threshold of 0.5. The area under the ROC curve (AUC) is a measure of the discriminatory power of a binary classifier and ranges from 0 to 100 (0 to 1 multiplied by 100). SDM predictions with AUC scores <50 have relatively equivalent true and false positive rates and fail as a classifier. AUC scores >50 indicate classifiers that more frequently predict occurrence accurately, but scores above 90 identify a strong classifier (Fielding and Bell 1997; Lobo et al. 2008).

To identify how evolutionary processes inform species environmental response, we compared response curve estimates for each species when estimated in *BePhyNE* with and without occurrence data using a leave-one-out approach in two steps. First, we fitted *BePhyNE* to the training dataset 1000 times, leaving out the occurrence data from one species each time. The recovered response curve parameter estimates for each data-dropped species were only informed by the occurrence of other species through the mvBM process. Second, we predicted occurrence patterns over the validation dataset and calculated AUC scores for data-dropped species range predictions. For each species, we compared these data-dropped AUC scores to the full data AUC scores calculated earlier.

Because we were particularly interested in understanding whether phylogenetic information was influencing estimates of ecological niches, we sought to compare the ecological niche estimates obtained from the evolutionary process and those using each species, even when occurrence data for that species is lacking. To evaluate this we measured the Mahalanobis distance of the optima and breadth for temperature and precipitation between the missing data posterior and each posterior sample from the data-informed posterior iteratively for each species. Thus, for each species we obtained a distribution of Mahalanobis distances between the data-informed posterior and the missing data posterior distribution. The missing data posteriors varied in their specificity, depending on phylogenetic position and the amount of data from related species. Thus, we repeated the Mahalanobis distance calculation, but measured it between the target species’ missing-data posterior and every other Plethodontid species’ data-informed posterior distribution. We then pooled the resulting distribution to identify the distance between the target species’ missing-data posterior and that from all other salamander species, thereby generating a null distribution. We expect the distances between the target species and the missing-data posterior to be on average closer to the target species than other salamander species randomly sampled across the phylogeny. We therefore measured the quantiles of the target species distances (*P_target_*) within the null distribution, and tested for deviations from uniformity with a KS-test. While predictions are expected to overall be improved by phylogeny, some taxa and clades may deviate from Brownian Motion niche evolution. For example, our model assumptions may be broken in clades that speciate by ecological speciation and character displacement (or in clades with an abundance of cryptic species). Thus, prediction accuracy may itself be distributed phylogenetically. We therefore estimated the phylogenetic signal in the median values of *P_target_* using Blomberg’s *K*, and tested for statistical significance using a randomization test as implemented in the function *phylosig* in the R package *phytools*. We interpreted significant phylogenetic signal in this statistic as an indicator of phylogenetic structure in the deviations from our Brownian Motion assumptions of evolution.

## IV. Results

### 1. Simulation Tests

Our simulations suggest that we can reliably recover parameter estimates under model assumptions, but not without some biases (Supplementary Fig. S1). Though true parameters roughly resembled a uniform distribution across MCMC posteriors, they did not fit closely enough to pass the Cook (2006) test. Despite biases in the estimation, MCMC posterior medians frequently recovered the data-generating true values for all parameters with high accuracy and precision (Fig. 3). Posterior medians for niche response curve parameters were the most accurate and unbiased, with *θ* being the most accurate. Overall, MCMC posterior medians for most mvBM parameters were less accurate than species response curves. *A_ω_* and ***σ***_ω_ were slightly overestimated at low true values and underestimated at high true values. *R_COR_* estimation bias shared the same direction but greater magnitude and lower accuracy than other model parameters. Though ***σ*** and *R_COR_* estimation accuracy was lowest, they improved in accuracy as the size of the dataset increased (Fig. 4). Evolutionary parameters increased in precision with increasing phylogenetic sample size, while tip parameters were not affected, as these are primarily affected by the size of the species-level occurrence data.

**Figure 3.**
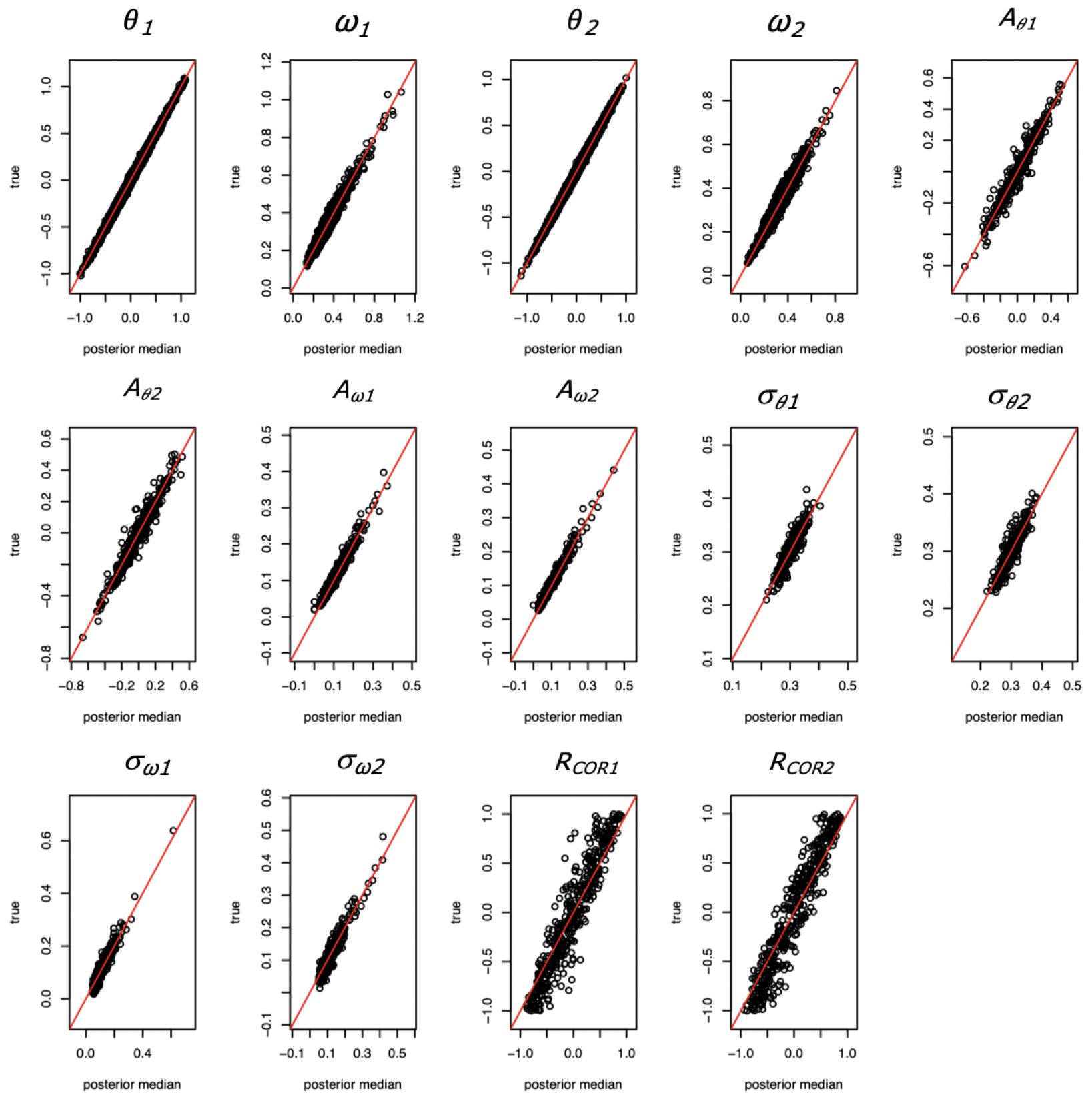
Estimated parameter performance. Estimated parameters (x-axis) are measured against “true” data generated parameters (y-axis) for two simulated environmental parameters (denoted 1 and 2). For tip parameters (*θ* and ω), plots show data for all species with each point indicating a single tip niche estimate.

**Figure 4.**
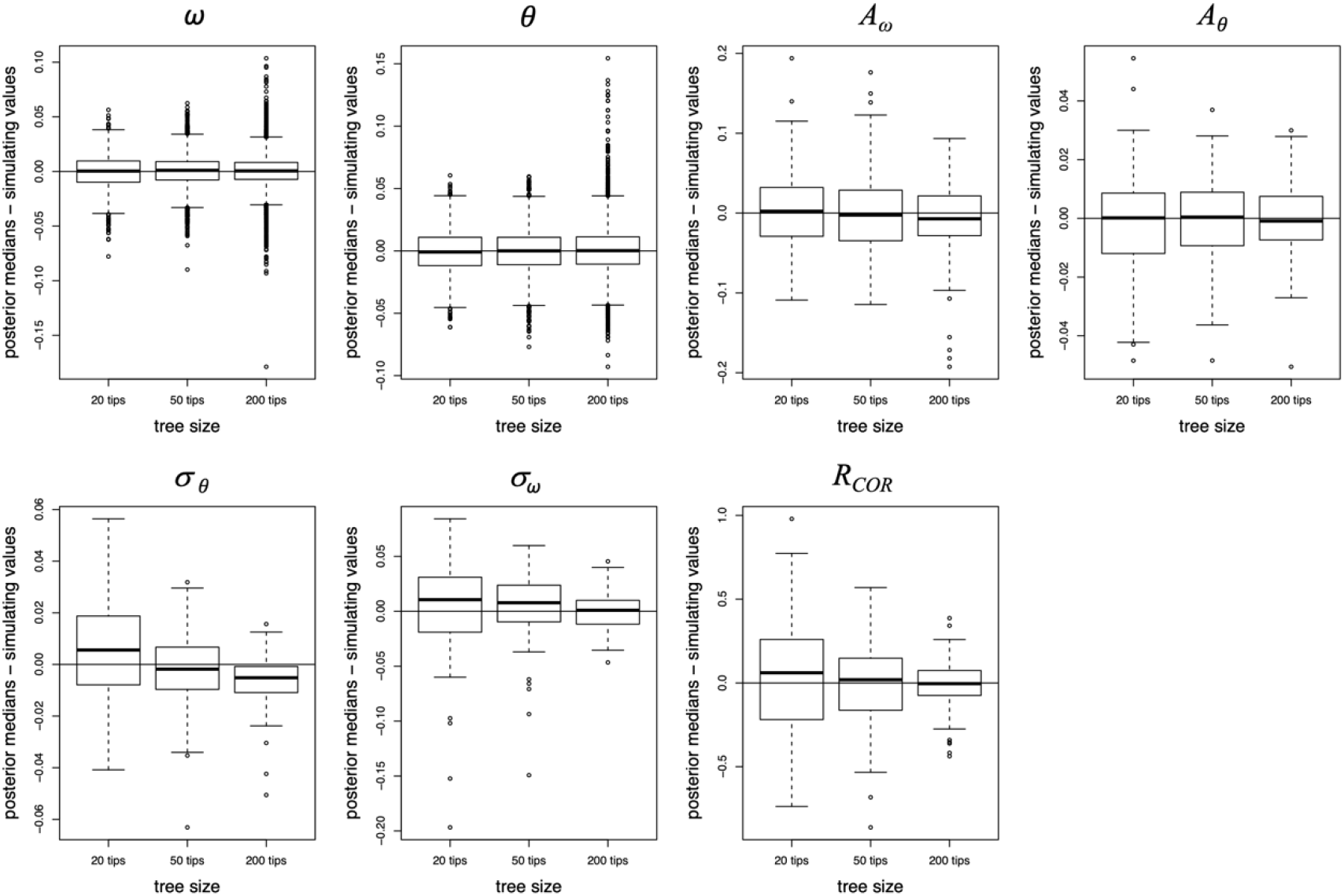
Parameter estimation accuracy, precision, and bias change with different tree sizes (20, 50, and 200 tips). Here, an overlap with 0 indicates higher accuracy.

### 2. Empirical Results: Eastern North American Plethodontidae

When fitted to the full dataset, we recovered similar estimates between *BePhyNE*, GLM and *BePhyNE* with dropped occurrence data. Closely-related taxa had more similar estimates in *BePhyNE* than in the GLM-only. For the mvBM model estimates (Table 1), posterior medians for *A_θ_* were 137.24 mm (95% HPD: 120.16–154.85 mm) for annual precipitation and 12.2°C (95% HPD: 9.63-14.76°C) for annual temperature. For *A_ω_*, posterior medians were 9.94 mm for annual precipitation (95% HPD: 6.82–13.81mm) and 1.70°C for annual temperature (95% HPD: 1.17–2.35°C). For *σ_θ_* posterior medians were 868.47 mm per million *years* for annual precipitation (95% HPD: 719.31-1,036.84 mm) and 12.04°C per million years for annual temperature (95% HPD: 10.05–14.29°C). For *σ_ω_*, posterior medians were 321.84 mm per million for annual precipitation (95% HPD: 235.01–427.25 mm) and 8.64°C per million for annual temperature (95% HPD: 6.71–11.09°C). The posterior distribution for annual precipitation correlation between *θ* and *ω* was largely above 0, with a posterior median of 0.10 (95% HPD: −0.01–0.21). Conversely, the posterior distribution for the same correlation for annual temperature was negative and below zero, with a posterior median of −0.15 (95% HPD: −0.25 – −0.05). Tip response curve estimates are clustered approximately by genus, with larger *ω* estimates in *Desmognathus* and *Eurycea* (Fig. 5).

**Figure 5.**
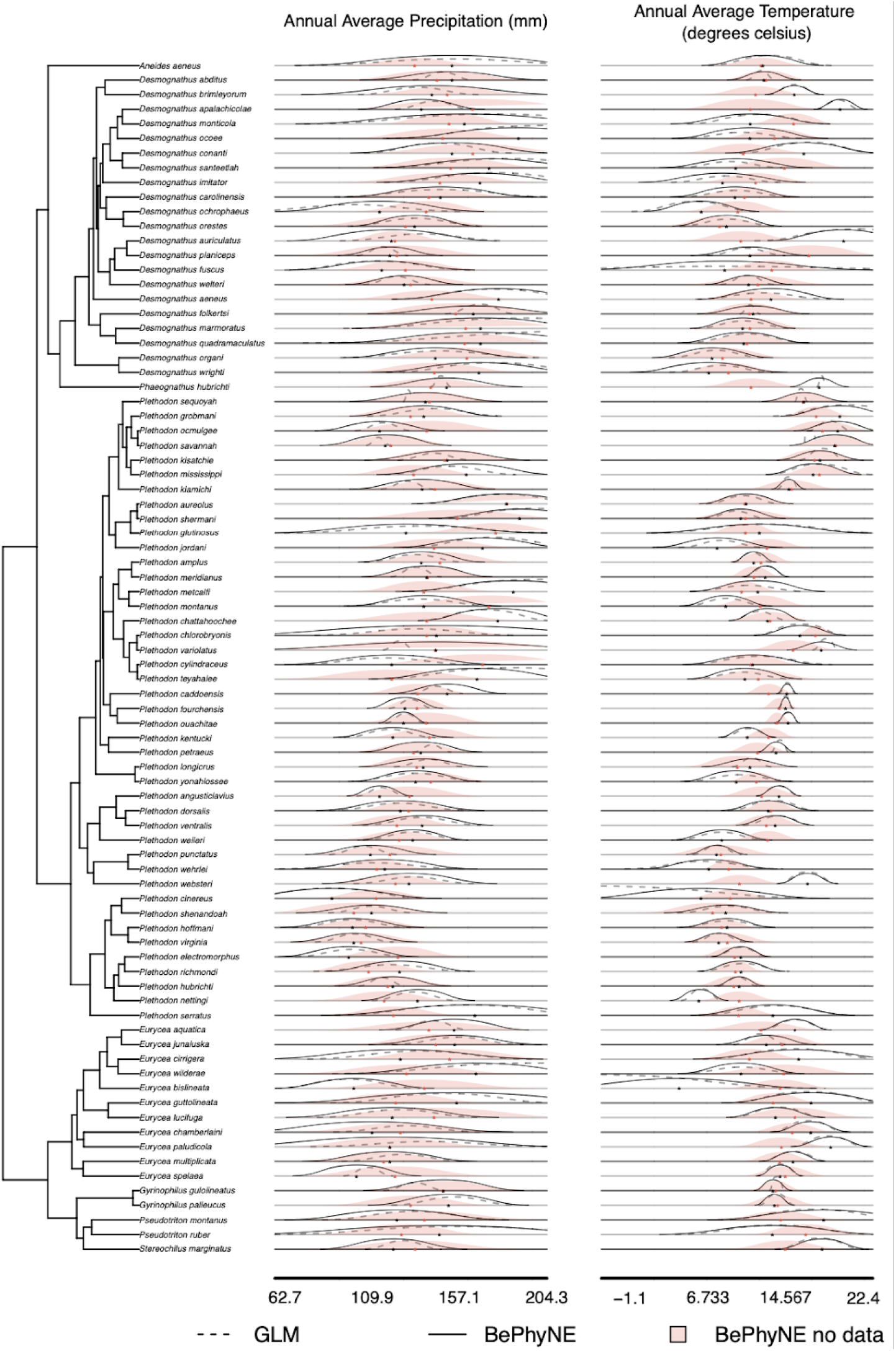
Response curve estimates mapped to phylogeny for the two climatic predictors, and for each of the three set-ups: *BePhyNE* (solid lines)*, BePhyNE no data* (estimation using the evolutionary model only; pink shadow), and traditional *GLM* (non-phylogenetic estimation of niche; dashed line).

Cross-validation supported the results of *BePhyNE* estimates for species niche response curves. AUC scores for prediction over the validation dataset varied across the family from 85-99 (Fig. 6). Predictions from *BePhyNE no data* generated slightly lower AUC scores for all species (82-99).

**Fig 6.**
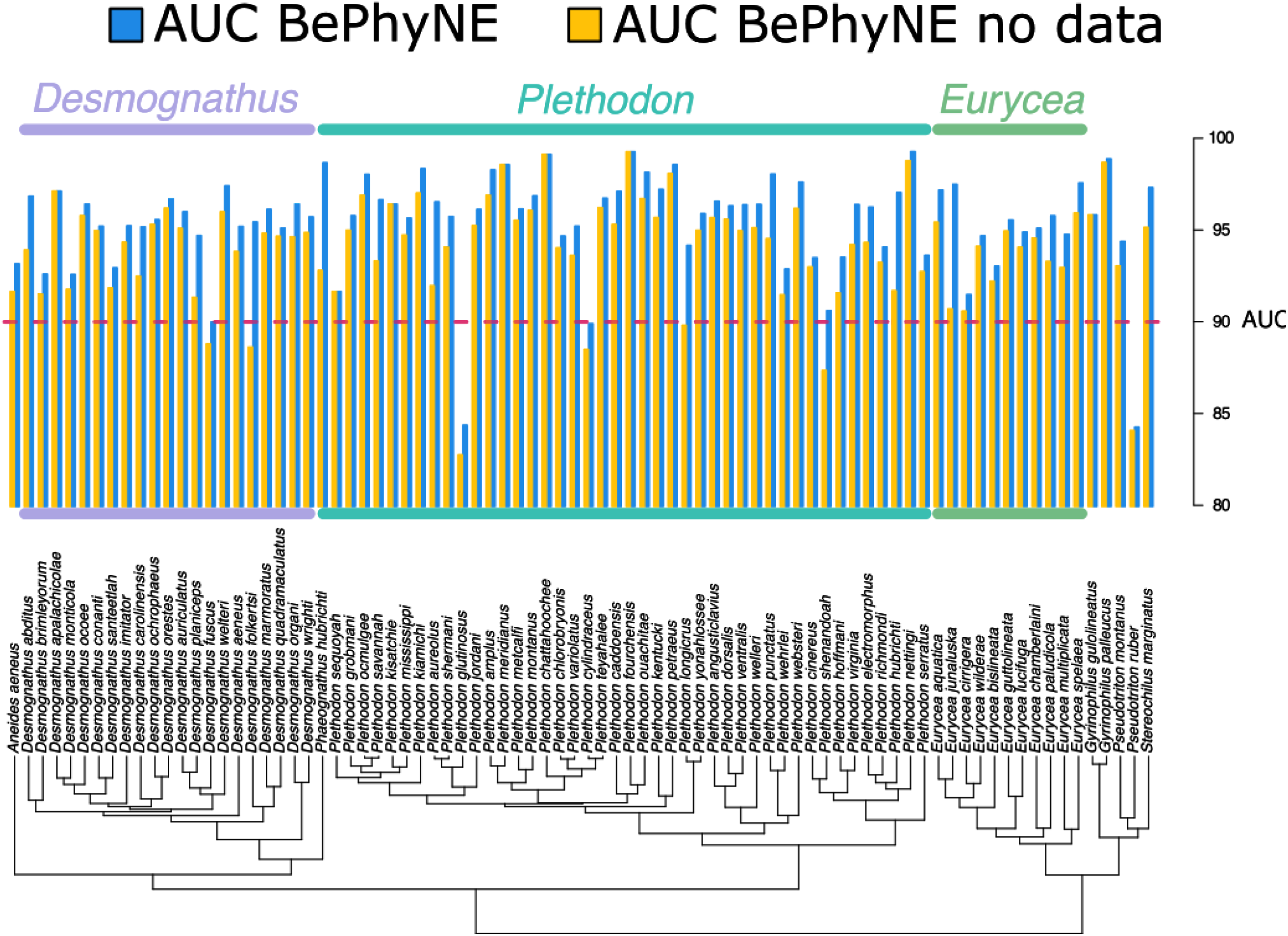
AUC scores for species niche estimates using *BePhyNE* (blue) and *BePhyNE* no data (yellow) setups for all Plethodontid species in the phylogeny

Results from our leave-one out cross-validation analysis demonstrate that the phylogeny does indeed inform estimates of ecological niche, even when species have no occurrences. In other words, *BePhyNE* can broadly inform estimates of a species’ ecological niche by leveraging information from related species and the evolutionary model. Posterior estimates of optima and breadth for temperature and precipitation estimated for species without data are all significantly associated with the posterior medians estimated with species occurrence data (Fig. 7). Furthermore, Mahalanobis distances between the multivariate niche from *BePhyNE* estimates lacking occurrence data and those with full occurrence data are much shorter than expected (Fig. 8). This is as evident by the significant left skew toward *P_target_* values near 0, which indicates that the distances are smaller than those measured against other randomly-sampled species of salamanders (Fig. 8). In addition, optimum (*Θ*) and breadth (*ω*) for both precipitation and temperature response curves show phylogenetic signal in their distance from the data-informed posterior distribution (*P _target_* has significant phylogenetic signal, Blomberg’s K = 0.28, p < 0.01; Figure 8). This indicates that there is phylogenetic signal in the prediction accuracy of salamander niches and may suggest phylogenetically-structured deviations from the Brownian Motion model of evolution. We further examined phylogenetic signal separately for temperature and precipitation response curves, and only found it in temperature curves (K = 0.31, p = 0.001) but not precipitation (K= 0.09, p = 0.95). This suggests that deviations from multivariate Brownian Motion may be more common in temperature, as might be expected if ecological speciation across temperature gradients drives divergence and results in significant displacements in the thermal niche relative to background rates, resulting in poor prediction from related species. Alternatively, these deviations may indicate portions of the tree with phylogenetic error and/or cryptic species.

**Figure 7.**
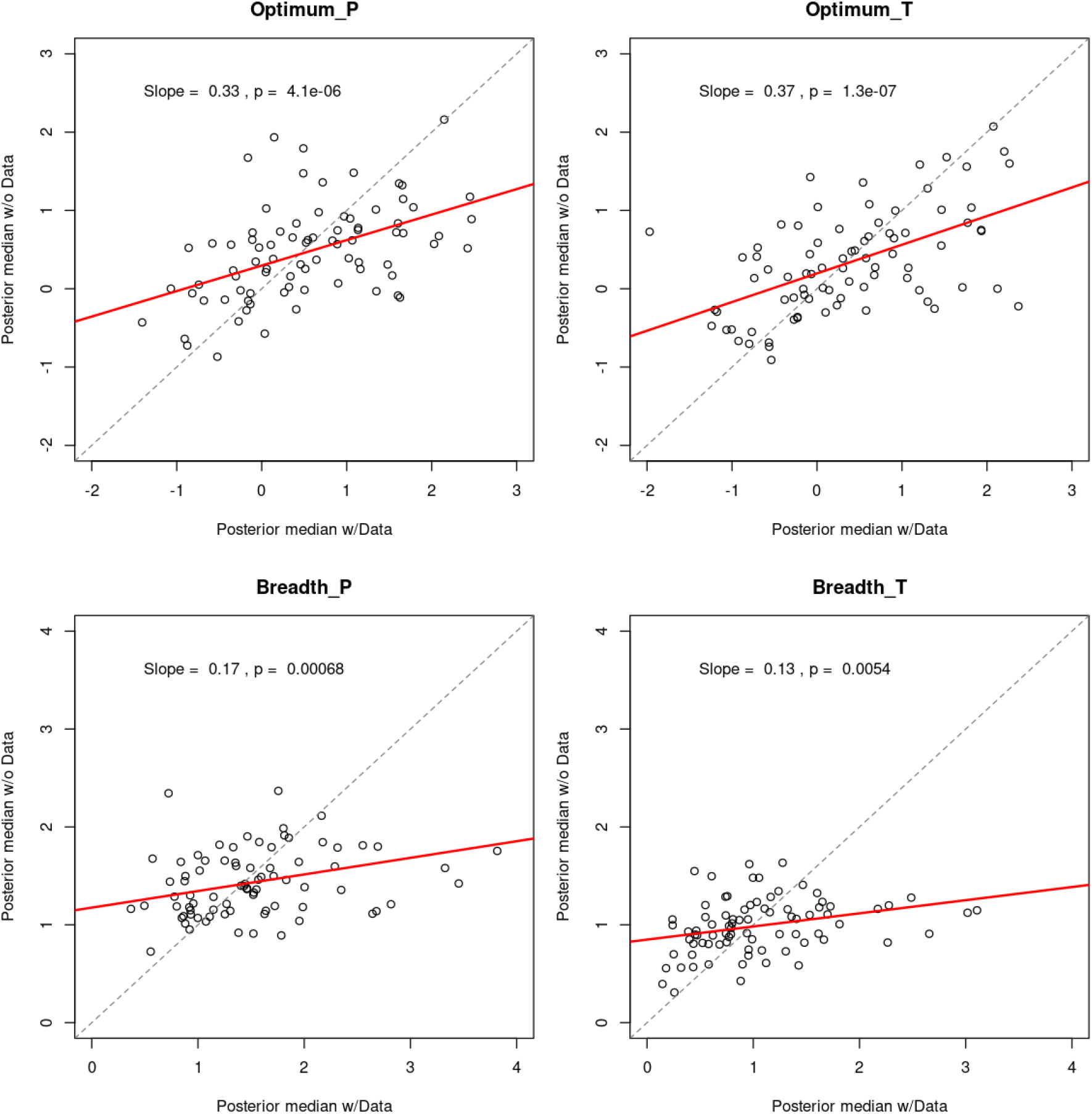
Relationship between posterior median estimates obtained using *BePhyNE* with species occurrence data (x) and *BePhyNE* no data (y), for A) precipitation *θ*, B) temperature *θ*, C) precipitation *ω*, and D) temperature *ω*. Red line indicates best-fìtting linear models, dotted gray lines indicate the expected pattern of a perfect correlation.

**Figure 8.**
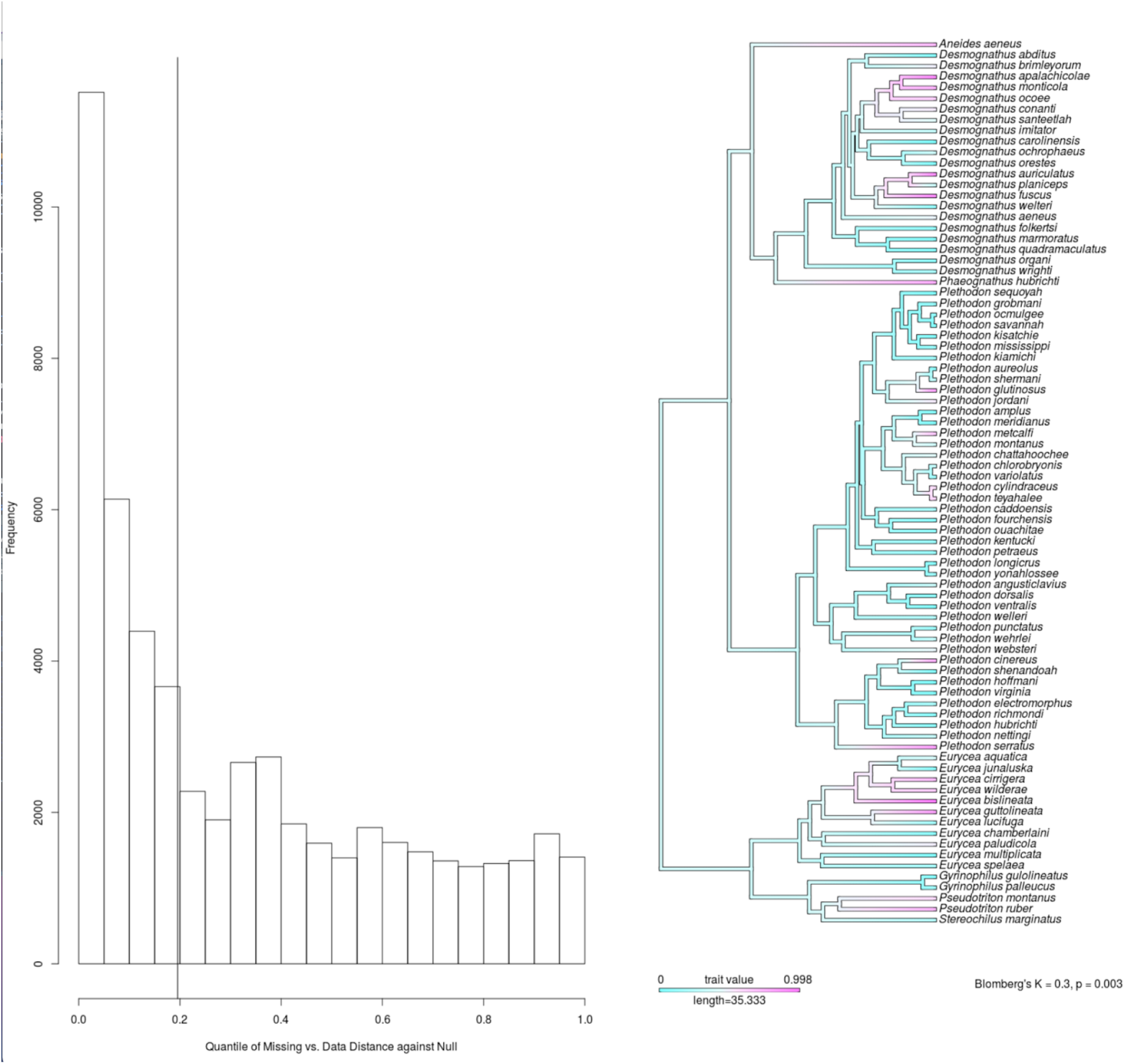
BePhyNE’s phylogenetic information improves ecological niche estimation in data-defìcient species A) Posterior quantiles of the Mahalanobis distances between the target species’ missing data posterior and the data-informed posterior within the randomized null distribution. Values skewed to the left indicate that the target species has a smaller distance to the missing-data posterior than species randomly sampled across the phylogeny. Vertical line indicates the average distance per species and its position in relation to the null distribution (0.22 quantile). This demonstrates that. B) Species median value for distance-quantile (*P_target_*) mapped on the phylogeny. Colors indicate distance between niches estimated using occurrence-rich and occurrence-poor setups. Blue: small distance; magenta: large distance. Significant Blomberg’s K indicates phylogenetic signal (K = 0.28, p < 0.01).

## V. Discussion

Here, we propose a novel method of phylogenetic niche estimation, *BePhyNE*, that jointly estimates species niches and the evolutionary process of niche evolution over the phylogeny. Previously, niche evolution was generally modeled using a two-step process where evolutionary models are fit to measured environmental means, or species specific ENMs (Lawing and Polly 2011; Lawing et al. 2016; Guillory and Brown 2021). However, such two-step estimates often carry considerable uncertainty from the original dataset and lack the ability to allow reciprocal information, with estimates of the evolutionary process model informing niche parameters for an individual species and vice-versa. Our approach further establishes a modeling framework to ask numerous questions of interest to evolutionary ecologists, such as: how is niche response *θ* and ω correlated along the phylogeny?, how will species evolve in response to climate change?, what is the environmental niche of rare and under-sampled species? By demonstrating the feasibility of joint modeling and parameter estimation, *BePhyNE* establishes a proof-of-concept modeling framework that demonstrates key improvements over the traditional two-step processes.

### 1. Joint Estimation of Species Niche and Evolutionary Process

*BePhyNE* is the first method to jointly estimate contemporary environmental response and BM processes in a Bayesian hierarchical framework. Our simulations showed that *BePhyNE* accurately estimates data-generating model parameters in most cases, but with some biases in posterior distributions, particularly for the evolution rate matrix parameters (***σ*** and *R_COR_*). However, even when posterior distributions were biased, posterior medians still closely approximated the data-generating model parameters. Our simulations also demonstrated model consistency, with larger phylogeny size improving model fits. Model performance may differ from our simulations under extreme evolutionary scenarios not included in the prior, including extreme instances of niche conservatism or heavily truncated realized niches across the clade. Our assumption of a random walk evolutionary process constrains niche divergence, resulting in more similar niche estimates in *BePhyNE* for closely related species than in GLM-only models (Fig. 5). This model behavior may lessen the impact of species-specific sampling biases in clades where the climatic niche is strongly shaped by macroevolutionary processes. We see this as being particularly useful for studying clades with large evolutionary divergences in niche due to abiotic dispersal barriers, as we expect closely-related species may have more similar fundamental environmental niches than expected from simply sampling realized occurrence patterns.

### 2. Practical Applications of *BePhyNE*

*BePhyNE* is unique among niche modeling approaches in that it can generate a model of environmental response over phylogeny that predicts distributions of poorly-understood species with limited to no occurrence data. Furthermore, the same model can be used to hind- and forecast spatial ranges. Using leave-one-out cross-validation, we demonstrated that many species in our empirical salamander example can have reasonable niche estimates even without occurrence data (Figs. 6–8).

Although very effective, *BePhyNE* can underperform in several conditions. First, this may be the case when the species that lack data are wide-ranging geographic generalists (e.g., *Eurycea bislineata, Plethodon cinereus*, and *Plethodon glutinosus*). Without occurrence data, generalists response curves tend to be pulled toward unrealistically narrower breadths to resemble the range restricted specialists to which they are often related. This said, climatic generalists are often the ones for which abundant data is available, meaning that this situation is less likely to occur in realistic conditions. Second, the ability to predict species’ niches using *BePhyNE* without occurrence data may vary across the phylogeny due to clade-specific shifts in evolutionary history, as suggested from the phylogenetic signal in the distance quantiles (Fig. 8). Such information may be leveraged to understand processes of niche divergence and diversification across the phylogeny. For example, our model had the lowest predictive power for genera *Desmognathus* and *Eurycea*. Because both genera are stream-adjacent specialists, they may be more buffered from macro-climatic change than terrestrial taxa due to their more thermally stable microhabitats in fast-moving streams (Steel et al. 2017; Shah et al. 2020), thus increasing the lag between macro-climatic change and niche evolution (Farallo et al. 2020). To account for variation in micro- and macro-climatic tracking, it is likely that using micro-climatic data whenever available will improve connection to physiological tolerances, as well as using macro-climatic characters that better characterize climate, including degree days and temperature ranges (Title and Bemmels 2018). Finally, because it relies on the phylogenetic relationships, the quality of the phylogeny may also impact the quality of the inferences. In our case, the taxonomic relationships of many Plethodontids (Camp and Wooten 2016; Pyron et al. 2022) are poorly understood and may bias our model and reduce predictive power. Resolving the biodiversity in the clade could potentially impact both species niche and distribution predictions and niche evolution estimates (Molbo et al. 2003; Gill et al. 2016).

Given the high proportion of species whose conservation status is listed as data deficient across the tree of life (IUCN 2021), our approach may allow for leveraging available phylogenetic information to predict distributions and understanding the niches of poorly-sampled species of conservation concern. Currently, few reliable methods exist for estimating niches of species without occurrence data, except through inference with sister taxa (Kozak and Wiens 2006; Eaton et al. 2008; Warren et al. 2008; Anciães and Peterson 2009). Cross-validation support for species response estimates with missing data suggests our model can recover estimates for data-deficient clades with phylogenetically-conserved niches. However, we also note that there is substantial room for improvement and expansion of the *BePhyNE* modeling framework. MCMC mixing for tips with missing data was slow for tips with fewer occurrence points. Future work to expand on proposal mechanisms to improve mixing will also facilitate niche estimation.

### 3. Inferring Environmental Response Through Time

In addition to estimating ranges for poorly-sampled taxa, our approach builds on previous methods used for estimation of niche evolutionary history and ancestral distributions (Pelletier et al. 2015; Guillory and Brown 2021). Often, models of ancestral distribution are generated by fitting paleoclimatic data to extant species niches, or two-step estimated response curves estimated on extant species and occurrence data. *BePhyNE’s* model of environmental response enables an ancestral range to be predicted over phylogeny directly from response curves. Given a time calibrated phylogeny, ancestral niche estimates can be approximated to specific periods with available paleoclimate to assess range shifts for species and clades through time. Though individual ancestral states estimates are unreliable (Schluter et al. 1997; Oakley and Cunningham 2000; Ekman et al. 2008), general patterns across ancestral states may still be informative and comparable – especially when considered with uncertainty across the full posterior distribution of reconstructed response curves. Although not tested here, expansion of this approach may facilitate the identification of historic processes shaping range.

Given our approach models a continuous process of niche evolution, we can play the evolutionary “tape” forward just as we can play it backwards (Gould 1990). The mvBM estimates of evolutionary rate in niche, as captured by *R*, can be used to generate possible stochastic future niches that could be expected for a species. Niche evolution may occur rapidly (Evans et al. 2009; Ogburn and Edwards 2015), enough so that forecasting under future climate models can identify scenarios where species may persist or go extinct.

### 4. Expanding the *BePhyNE* modeling framework

While a random walk is an inherently simplistic model of evolution for complex niche histories, by connecting niche estimation directly to the large family of continuous trait evolutionary models, we enable a wide range of possible modifications to *BePhyNE* that are straightforward to implement. For example, the *BePhyNE* modeling framework could be expanded to Ornstein-Uhlenbeck models that capture adaptation toward an optimal state (Kozak et al. 2005, 2006; Camp and Wooten 2016; Weaver et al. 2020), adaptive shifts (Butler and King 2004), relaxation and canalization of niche constraints (Beaulieu et al. 2012), competition (Drury et al. 2016), accelerating or decelerating niche evolution (Blomberg et al. 2003), and Levy Process models that allow for large “jumps” in niche evolution (Landis et al. 2013; Uyeda and Harmon 2014). Furthermore, the Bayesian implementation of *BePhyNE* enables the use of informative priors. Besides root estimates and evolutionary rates, fossil occurrences can be integrated as node priors for continuous trait evolution, improving estimation by calibrating responses based on paleontological data (Slater et al. 2012, Lawing et al. 2016, Rivera et al. 2020).

*BePhyNE’*s response curves could also be connected to species-level estimates of environmental tolerances, such as the thermal performance curves estimates in many physiological studies (Angilletta and Angilletta 2009; Drake 2015; Jiménez et al. 2019; Soberón and Peterson 2020; Jiménez and Soberón 2022). Exactly how our estimates, which incorporate abiotic and biotic impacts on niche width and optimum, are connected to the physiological tolerances of species is an open question. Such performance curves may often deviate from our symmetric gaussian curves, showing skewness (Austin et al. 1994; Austin 2002; Lawesson and Oksanen 2002), and some curves may be unbounded directional preferences (Bio et al. 1998; Lambert et al. 2011). Further incorporating these possibilities into our framework and connecting them with physiological data may provide new insights into the connection between micro-scale physiological tolerances and macroecological niche evolution.

### 5. Plethodontid Salamander Niche Evolution

We applied *BePhyNE* to a radiation of Plethodontid salamanders and evaluated its performance with AUC and leave-one-out cross-validation. This clade is of particular interest due to its dynamic niche evolutionary history and conservation relevance (Kozak and Wiens 2006, 2010a, 2010b; Milanovich et al. 2010; Kozak 2017). Plethodontidae salamanders are lungless and respire through their skin, thus requiring wet and cool conditions to survive. Across their Eastern North American range, Plethodontidae are largely divided between widely-distributed lowland species and range-restricted species endemic to high-elevation localities in the south Appalachian Mountains. Though not fully understood, historic patterns of climate change and niche specialization to cool and wet habitats likely resulted in current day Plethodontidae ranges (Kozak 2017). Increasingly drier and warmer conditions restricted the range of climatic specialists to high elevation zones where lineages then accumulated (Kozak and Wiens 2006, 2010b; Shen et al. 2016). Temperature particularly, is likely to constrain the distributions of many of the high elevation endemics, especially on their lower elevational limits within the warmer Southeastern U.S.A. (Kozak and Wiens 2006; Arif et al. 2007; Gifford and Kozak 2012). Agreeing with this biogeographic scenario, our phylogenetic niche model supports a pattern of greater thermal constraint, with wider relative *ω* in precipitation than temperature across most species. This is of conservation relevance, since with rising temperatures due to climate change, many montane specialist species will be trapped in a shrinking habitat, with predicted ranges shifting to higher elevations until their climatic optima disappear and the species becomes extirpated (Vieites et al. 2009; Milanovich et al. 2010; Gifford and Kozak 2012; Moskwik 2014).

We also identify negative evolutionary correlations in thermal *ω* and *θ*, and positive correlations in precipitation *ω* and *θ*, suggesting species that occur within the warmest and driest parts of Eastern North America have the narrowest niche breadths. This pattern between precipitation and thermal niche breadth is supported by several previous studies across Amphibia (Bonetti and Wiens 2014), and some lizard clades (Wiens et al. 2013; Lin and Wiens 2017; Mengchao et al. 2019), suggesting a more general macroecological pattern of niche specialization in niche evolution (Sexton et al. 2017; Carscadden et al. 2020). However, variation in Plethodontid evolutionary patterns is possible, as subclades differ in habitat and natural history (Kozak et al. 2005, 2006; Camp and Wooten 2016; Weaver et al. 2020). Indeed, several species with the most aquatic habitats, including *Desmognathus* and *Eurycea* have the widest *ω* estimates for both predictors. There are many potential reasons for these patterns, including greater resilience to ambient air temperatures compared to the more terrestrial species (Myers and Adams 2008). Though a small sample in our study, this pattern of wide niches in semi-aquatic salamanders is also notable in *Pseudotriton*, a largely semi-aquatic genus of salamander with nearly the widest niches (Bruce 1975; Marvin 2003).

The differential histories of many Plethodontid subclades may explain the differences in niche breadth and optimum estimates between *BePhyNE* and the GLM-only model (Fig. 5). The cryptic and putatively “non-adaptive radiation” (Kozak et al. 2006) of *Plethodon glutinosus* exhibits the most divergent response curve patterns between models. *BePhyNE* niche breadth estimates were wider, resembling closely-related taxa more than the narrower GLM-only niche breadth estimates. This pattern supports previous research suggesting that species within the complex are ecologically similar. Over a history of climatic shifts driving range expansions and contractions (Webb III and Bartlein 1992; Jansson and Dynesius 2002), the clade diverged allopatrically and rapidly accumulated ecologically-conserved lineages geographically constrained to a subset of their realized distribution by competition (Losos and Glor 2003; Kozak et al. 2006). This is in contrast to the similar niche curves recovered by *BePhyNE* and the GLM-only model in *Desmognathus*. Many *Desmognathus* species diverged early in the clades’ history to adapt to numerous microhabitats, stably co-occurred through multiple climatic cycles to the present (Kozak et al. 2005). This historic pattern of community assembly has been maintained through interspecific competition and predation (Krzysik 1979; Hairston 1980; Keen 1982; Hairston, 1986). The overlapping niche estimates among species is expected, as biotically-structured and sympatric clades should exhibit little divergence in macro-climatic response (Ackerly et al. 2006; Li et al. 2018). We also observe similar niche curves recovered by *BePhyNE* and GLM-only models for the *Plethodon cinereus* complex, a clade of wide-ranging generalists and hyper-endemics. Many southern hyper-endemic specialists are range-restricted due to a history of environmental shifts restricting the availability of ideal microhabitats to specific mid-elevational bands. While some species may be climatically-restricted from other ideal habitats (Shen et al. 2016; Kozak 2017), others may be outcompeted by more widespread generalists (Jaeger 1971, 1972, 1974; Myers and Adams 2008; Lyons et al. 2016).

Though our phylogenetic niche model pulls species response away from realized occurrences and towards a more phylogenetically-conserved value, we emphasize we are not directly modeling the fundamental niche. Our model can reveal which species in a clade have particularly truncated niches relative to other members of a clade, but it would be unsurprising if some Plethodontid clades have conserved truncations due to prevalent dispersal or biotic interactions (Soberon and Peterson 2005; Soberon and Nakamura 2009). In such cases, we would not expect responses to differ between *BePhyNE* and non-phylogenetically informed SDMs. One way to reduce the risk of clade-wide niche truncation is utilizing physiological experiments, such as critical thermal maximum and minimum trials, to inform species *θ* and ω when possible. Integrating this information into the niche priors could allow testing the extent to which species responses estimated from occurrence data are pulled toward the fundamental niches of phylogenetically-related species.

### 6. Conclusions

We developed, implemented and tested a novel model for estimating ecological niches jointly for all species across a phylogeny by linking GLM estimation with an evolutionary process model for the underlying species response curves. We demonstrate effective estimation of niche and evolutionary parameters from simulated and empirical datasets. In addition, our framework provides a proof-of-concept for a new class of approaches for studying niche evolution, including improved methods for estimating niche for data-deficient species, historical reconstructions, future predictions under climate change, and evaluation of niche evolutionary processes across the tree of life. Our approach establishes a framework for leveraging the rapidly growing availability of biodiversity data and molecular phylogenies to make robust eco-evolutionary predictions and assessments of species’ niche and distributions in a rapidly-changing world.

## Acknowledgements

We would like to thank Daniel Caetano, Joel McGlothlin, Emmanuel Frimpong, Nic Bone, Bailey Howell, Fabio Machado, Diego Sasso Porto, and others for feedback and comments on early versions of this manuscript.

## SUPPLEMENTAL INFORMATION

### Supplement 1: Algebraic transformation of beta coefficients to niche characters

We consider the GLM equation used in phylogenetic niche modeling of measured predictor variables *X*, such as temperature or precipitation, for a set of presence/absence observations *Y_i_* for species *i*. A GLM can be used to estimate the relationship between the predictor and the probability of occurrence *P_i_* such that:

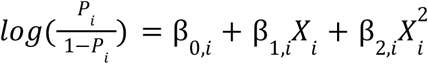

The observations *Y_i_* are Bernoulli distributed with probability *P_i_*. However, it is easy to choose parameters for β values that result in a preference/tolerance curve (hereafter, simply the preference curve) that is biologically unrealistic (for example, a species may not have an intermediate optimum for *X*, but rather an intermediate valley). Furthermore, the interdependence of these parameters in the equation means that they are difficult to interpret. To remedy this, we reparameterize the model and constrain the preference curve to correspond to a unimodal distribution around an intermediate optimum θ. The width of this curve around the optimum is given by ω, and the height of the curve at the optimum is given by τ. These have biologically interpretable values upon which we can set meaningful constraints.

To solve for θ in terms of β values, we take the derivative *dP/dX* after solving Eq. [eq1] for *P_i_*, and find the value of *X* where *dP/dX* = 0 (i.e. the maximum). We will, for now, drop the *i* subscripts designating species *i*. Rearranging Eq. [eq1]:

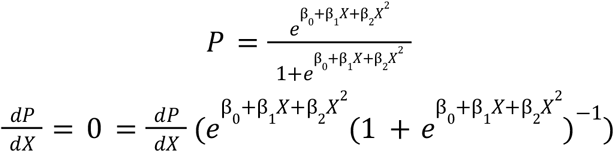

Using the product rule, this results in:

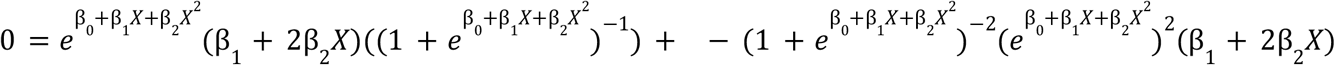

which simplifies to:

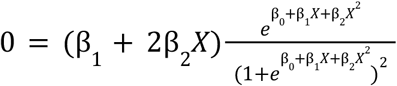

Thus, we find the optimum as:

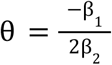

The height of the curve, τ, is simply Eq. [eq1] evaluated at θ.

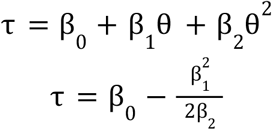

The width of the curve corresponds to half the distance between the two points on the curve where the probability of occurrence falls below a certain fixed value ν. In other words,

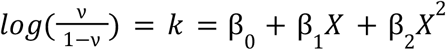

The width as we have defined it can be determined by taking half the distance between the two solutions to the quadratic formula, which is:

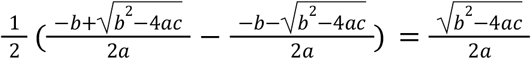

In our equation, this results in:

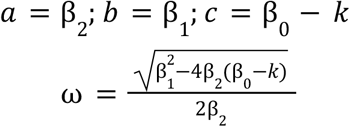

This model is of course subject to a number of constraints that we will impose. For example, τ > v, meaning that the occurrence probability at the optimum must be greater than the cutoff for measuring the width. Furthermore, β_1_ >0, β_2_ < 0, and β_0_ < 0 (not sure if these are exact).

Back transformation

To obtain values of β = [β_0_, β_1_, β_2_] from Θ = [θ, ω, τ], we solve the system of equations Eqs. [theta], [tau], [omega], and obtain:

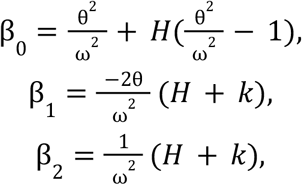

where 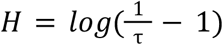 and 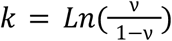

### Supplement 2: Cook (2006) test

Under the Cook (2006) test, if our model is accurate and precise, the true values will be uniformly distributed in the quantiles of the estimated posterior distributions (Cook et al. 2006). To test this, we measured the quantiles of the simulated parameter values from the empirical cumulative distribution (ECD) function of the MCMC posteriors. We expect that over a large number of simulations (>500) the “true” parameter values will be uniformly distributed over the ECD.

Our results show that our model does not pass the Cook (2006) test. The most uniformly-distributed parameters were the niche response optima *θ*. However, the MCMC posteriors did not recover the true parameters more often than expected, creating a sharp U-shape at the tails of the distribution. Tip response curve breadths *ω* MCMC posteriors did not contain the true values in the upper tail of the distribution as often as expected. *R_COR_* matrix MCMC posteriors were approximately uniformly distributed. The MCMC posteriors for the mvBM evolutionary rates ***σ*** were the most difficult to recover, with ***σ***_θ_ skewed towards underestimation, and ***σ***_ω_ skewed towards overestimation.

**Figure 1:**
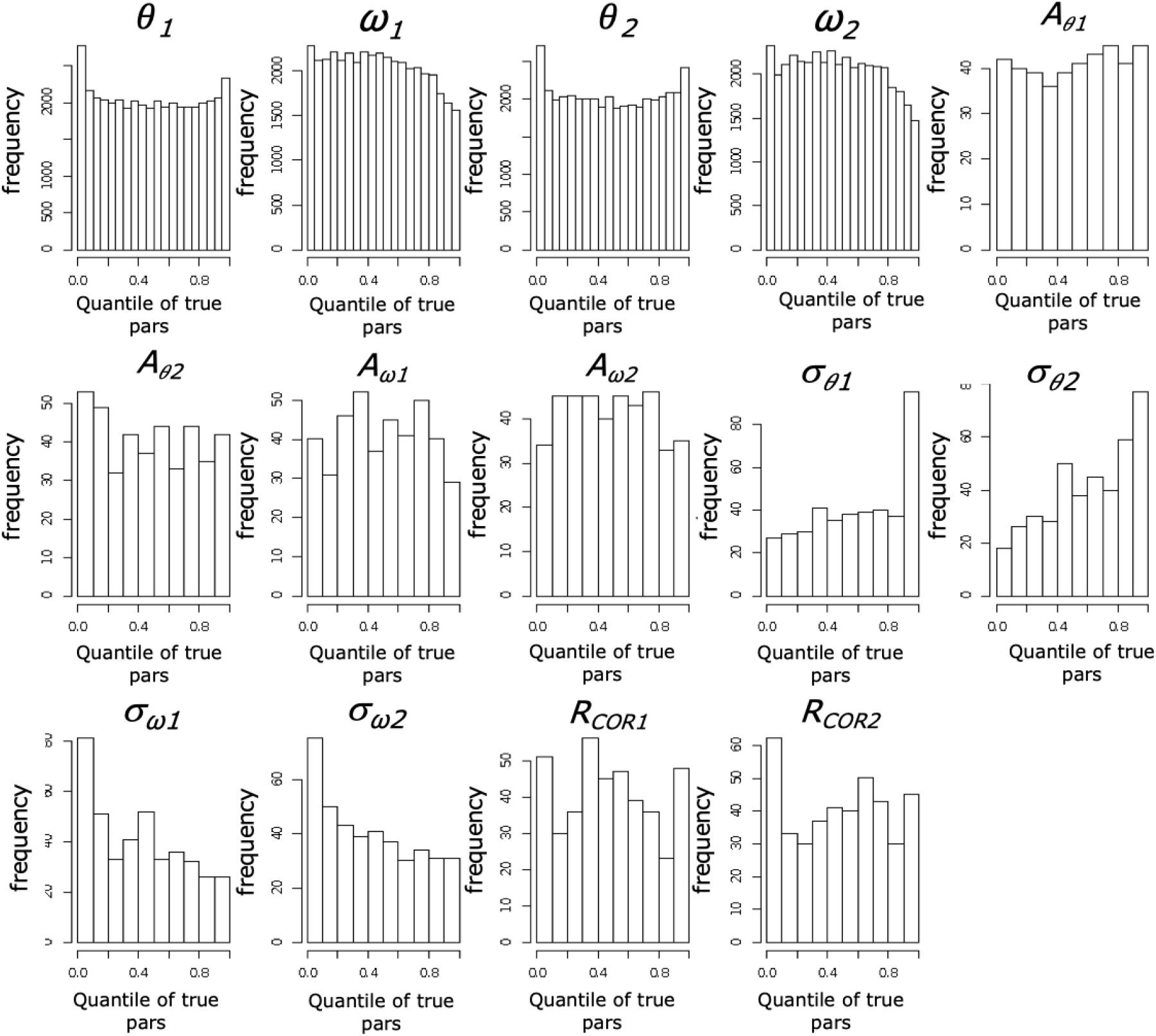
Empirical cumulative distribution plots for all parameters estimated in simulation analyses. We simulated and estimated parameters from 500 datasets sampled from the prior using 1 million generation MCMC at 10% burnin. For each dataset, we identified where “true” data generating parameter values fit on the empirical cumulative distributions, and plotted all 500 fits for each parameter as a histogram. Distributions should follow a uniform distribution if the model estimates parameters without bias and if model convergence is achieved. Our results indicate some parameters either have difficulty achieving convergence, or are subject to estimation bias. Particularly, the ***σ*** values for the multivariate Brownian Motion process are underestimated for the niche *θ*, and overestimated for niche ω.

